# Uncovering differences in rye and wheat degradation by human gut microbiota applying a quantitative multi-metaOmics *in vitro* approach

**DOI:** 10.1101/2025.06.06.658233

**Authors:** Clara Berenike Hartung, Alexandra Kelder, Sabrina Woltemate, Robert Geffers, Andreas von Felde, Knud Erik Bach Knudsen, Christian Visscher, Marius Vital

**Affiliations:** Institute for Medical Microbiology and Hospital Epidemiology, Hannover Medical School, Hannover, Germany; Institute for Animal Nutrition, University of Veterinary Medicine Hannover, Foundation, Hannover, Germany; Genome Analytics Research Group, Helmholtz Centre for Infection Research, Braunschweig, Germany; KWS Lochow GmbH, Bergen, Germany; Department of Animal and Veterinary Sciences, Aarhus University, Tjele, Denmark; German Center for Infection Research (DZIF), Hannover-Braunschweig, Germany

**Keywords:** gut microbiota, diet, wheat, rye, SCFA, propionate, butyrate, acetate, metagenomics, metatranscriptomics

## Abstract

While wheat is the most common grain used in bread-making worldwide, rye is popular in many European countries too. Rye is associated with several health benefits, which is attributed to its comparatively higher fiber content (primarily fructans and arabinoxylans) that promote production of short chain fatty acids (SCFA) by gut microbiota, in particular butyrate. Intervention studies revealed bacterial alterations upon rye administration, however, the detailed mechanisms involved in its degradation are not understood.

We grew fecal communities (n=20) on pre-digested rye and wheat, respectively, demonstrating that rye was yielding higher cell and SCFA concentrations in almost all samples along with distinct abundances of many taxa. A multi metaOmics (metagenomics/metatranscriptomics) approach (n=5 donors) showed higher bacterial growth rates for most taxa on rye compared to wheat. The higher growth rate of rye was accompanied by increased expression of genes involved in growth and energy generation suggesting higher carbon substrate accessibility. The carbohydrate active enzyme repertoire was greatly distinct between communities growing on the two substrates with several specific glycoside hydrolases increasingly expressed in rye containing cultures. *Agathobacter faecis* was revealed as the key butyrogenic species for rye degradation and its expression pattern based on metagenome assembled genomes showed adaptation to growth on rye via expression of genes involved in arabinoxylan degradation and fructose (major monomer of fructans) uptake.

Our study verifies higher SCFA production from rye over wheat and gives detailed insights into molecular mechanisms involved. It supports that the observed health benefits of rye are mediated by gut microbiota.

## Introduction

Understanding how diet governs gut microbiota composition and function is a major research effort today. A top priority is to unravel factors promoting the production of short chain fatty acids (SCFA) that are bacterial fermentation end-products and a hallmark of beneficial microbiota. SCFA provide epithelial integrity by feeding enterocytes and promoting mucus production ^1^, they have anti-inflammatory properties and, upon entering the circulation, exert beneficial effects on various peripheral sites such as the liver and the brain ^2^. The main SCFA are acetate, butyrate and propionate. Acetate is produced by the vast majority of gut bacteria, whereas butyrate and propionate are only formed by certain members that generally form separate groups ^3,4^. SCFA deficiencies have been shown to play a role in various emerging diseases including obesity, diabetes type 2 and cardiovascular disease among others^5,6^. Furthermore, SCFA sustain bacterial community homeostasis and provide colonization resistance against harmful bacteria by lowering the intestinal pH as well as by direct actions against pathogens ^7,8^.

Investigations on the diet - gut microbiota axis are performed on various levels involving studies focusing on specific substrates (e.g. prebiotics), individual food components or entire dietary regimes. Inulin and resistant starches are prominent examples for prebiotics and we recently performed a meta-analysis substantiating the common view that those compounds indeed stimulate the growth of specific SCFA-producing taxa, in particular certain butyrate-producers, where the community composition pre-intervention was uncovered to play a crucial role ^9^. Adherence to certain diets enriched in fibers, such as the Mediterranean diet, is another prominent line shown to increase SCFA synthesis ^10^. Unraveling the role of specific food components is in-between and aims to understand the impact of specific dietary components on gut microbiota function. Several foods that contain high fiber concentrations, such as legumes, fruits and berries among others, are known to promote SCFA production ^11-13^. Cereals are a major source of fibers too ^14^, where whole grains have been shown to promote higher SCFA formation over refined grains ^15,16^. The grain type is important as well. While wheat is the worldwide leading cereal by far, rye is an important bread cereal too, especially in central as well as eastern European countries and in Scandinavia ^17^. Due to its unique carbohydrate composition, rye contains a higher proportion of dietary fibers than wheat. It holds more fructans and soluble arabinoxylans than other cereals providing a high content of soluble non-starch polysaccharides (NSP) ^18,19^. Both fructans and arabinoxylans were shown to promote SCFA production ^20^, and especially for inulin, a specific type of fructan, a butyrogenic effect has been well established^9^. Recently, a dietary intervention study investigating the effect of a hypocaloric diet rich in either high fiber rye foods or refined wheat foods in a 12-week weight-loss trial demonstrated increased butyrate plasma levels along with higher relative abundances of the butyrate-producing bacterium *Agathobacter* in the former group ^21^. Furthermore, changes in butyrate and propionate were inversely correlated with changes in body fat percentage in the rye group, but not in individuals consuming wheat ^21^.

In summary, it is largely accepted that the positive effects of fibers on host health are (at least partly) mediated by gut microbiota-derived SCFA. However, the exact microbial mechanisms involved during fermentation of certain foods, specifically regarding its SCFA promoting compounds, are still largely in the dark hardly reaching beyond association studies that focused on the taxonomic level only. This is also true in the case of major cereals including wheat and rye. To this end, we performed various *in vitro* experiments on pre-digested rye and wheat applying multi metaOmics approaches to unravel microbial taxa and functions governing their degradation and subsequent SCFA formation.

## Results

### Investigating growth of fecal samples derived from 20 individuals on pre-digested rye and wheat

In order to obtain broad insights into communities grown on pre-digested rye and wheat we performed batch cultivations with fecal samples from 20 individuals. Major nutrient compositions of pre-digested substrates, compared with the native cereals, are shown in Table 1. During pre-digestion, the starch content in both cereals decreased notably. The protein content in wheat decreased as well, however, that was less pronounced in rye compared to wheat. The amount of fructan, soluble NSP, dietary fiber, arabinose and xylose was higher in rye than wheat with differences being greater in pre-digested cereals; overall, the share of those substrates also increased after pre-digestion.

**Table 1.** Starch, crude protein, non-starch polysaccharides (NSP) and lignin content (% of dry matter) of the native and pre-digested substrates according *(preprint coming soon*)

| % Dry matter |  | Wheat | Pre-digested wheat | Rye | Pre-digested rye |
| --- | --- | --- | --- | --- | --- |
| Starch |  | 63.1 | <30.0 | 64.0 | <30.0 |
| Crude protein |  | 14.3 | 12.4 | 8.12 | 10.4 |
| Enzymatic sugars |  |  |  |  |  |
| Glucose |  | 0.249 | 1.56 | 0.322 | 1.93 |
| Fructan |  | 0.142 | 1.15 | 2.19 | 6.82 |
| Non cellulosic polysaccharides (NCP) |  |  |  |  |  |
| total NCP | Soluble | 2.40 | 13.9 | 3.80 | 17.7 |
|  | Insoluble | 8.44 | 33.8 | 11.3 | 28.7 |
|  | Total | 10.8 | 47.8 | 15.1 | 46.4 |
| Arabinose | Soluble | 0.755 | 3.55 | 1.23 | 4.79 |
|  | Insoluble | 1.90 | 7.89 | 2.31 | 6.80 |
|  | Total | 2.66 | 11.4 | 3.54 | 11.6 |
| Xylose | Soluble | 1.08 | 6.79 | 1.84 | 7.53 |
|  | Insoluble | 3.14 | 12.6 | 3.63 | 9.87 |
|  | Total | 4.22 | 19.4 | 5.47 | 17.4 |
| Others |  |  |  |  |  |
| Cellulose |  | 1.52 | 8.18 | 0.754 | 3.86 |
| Lignin |  | 1.25 | 4.85 | 1.09 | 3.80 |
| Dietary Fiber | Soluble | 2.40 | 13.9 | 3.80 | 17.7 |
|  | Insoluble | 9.69 | 38.7 | 12.3 | 32.5 |
|  | Total | 12.1 | 52.6 | 16.1 | 50.2 |

Final cell concentrations reached after 24h were significantly higher in rye (1.42 × 10^9^ ± 0.25 × 10^9^ cells mL^-1^) than wheat (1.09 × 10^9^ ± 0.25 × 10^9^ cells mL^-1^) with all intra-individual comparison displaying more growth with the former substrate (difference of 3.30 × 10^8^ ± 1.55 × 10^8^ cells mL^-1^) (Figure 1a). Similar for SCFA, where all three SCFA reached higher concentrations in rye than wheat; acetate (rye: 11.12 ± 2.29 mM; wheat: 8.78 ± 2.56 mM; difference of 2.34 ± 1.13 mM), butyrate (rye: 2.88 ± 0.57 mM; wheat: 2.34 ± 0.46 mM; difference of 0.55 ± 0.29 mM) and propionate (rye: 2.35 ± 0.74 mM; wheat: 2.08 ± 0.63 mM; difference of 0.26 ± 0.23 mM) (Figure 1b,c); only one individual each showed lower concentrations on rye (#1 for acetate and butyrate; #17 for propionate). Ordination analyses of grown communities based on 16S rRNA gene data clustered samples of individual subjects together with all replicates showing tight associations and little substrate-specific signatures. However, substrate did significantly influence community composition based on PERMANOVA analysis (p <0.05) (Figure 1d). Based on absolute abundance data samples of the two substrate groups showed clear separation (Figure 1e). Furthermore, a multitude of taxa differed significantly in their relative abundances demonstrating that substrates stimulated growth of specific taxa. For instance, the genera *Agathobacter*, *Anaerostipes* and *Blautia_A* were enriched after growth with rye, while *Phocaeicola* and *Sarcina* were higher in wheat derived samples (Figure 1f; a detailed list of differentially abundant taxa is given in Table S1). Random Forest analyses could discriminate samples between the two groups at high accuracies (AUC of 0.97 (relative abundance) and AUC of 0.95 (absolute abundance) reflecting two and four false classifications, respectively) (Figure 1g). Communities grown on medium alone showed much lower cell concentrations (4.2 × 10^8^ ± 0.77 × 10^8^ cells mL^-1^) and greatly distinct compositions, with much higher concentrations of *Escherichia* and *Parabacteroides* among others (Figure S1a), as did incubations with only substrates (rye: 4.12 × 10^8^ ± 0.45 × 10^8^; wheat 3.50 × 10^8^ ± 0.15 × 10^8^ cells mL^-1^) that contained no fecal communities and were dominated by *Escherichia* and *Sarcina* (Figure S1b). For above analyses, data from one individual (#4) were omitted due to their stark compositional differences compared to all other samples (Figure S1b). Together, our results revealed clear differences in bacterial communities growing either on rye or wheat, respectively, where the former stimulated higher overall growth along with a specific community structure and increased SCFA concentrations.

**Figure 1.**
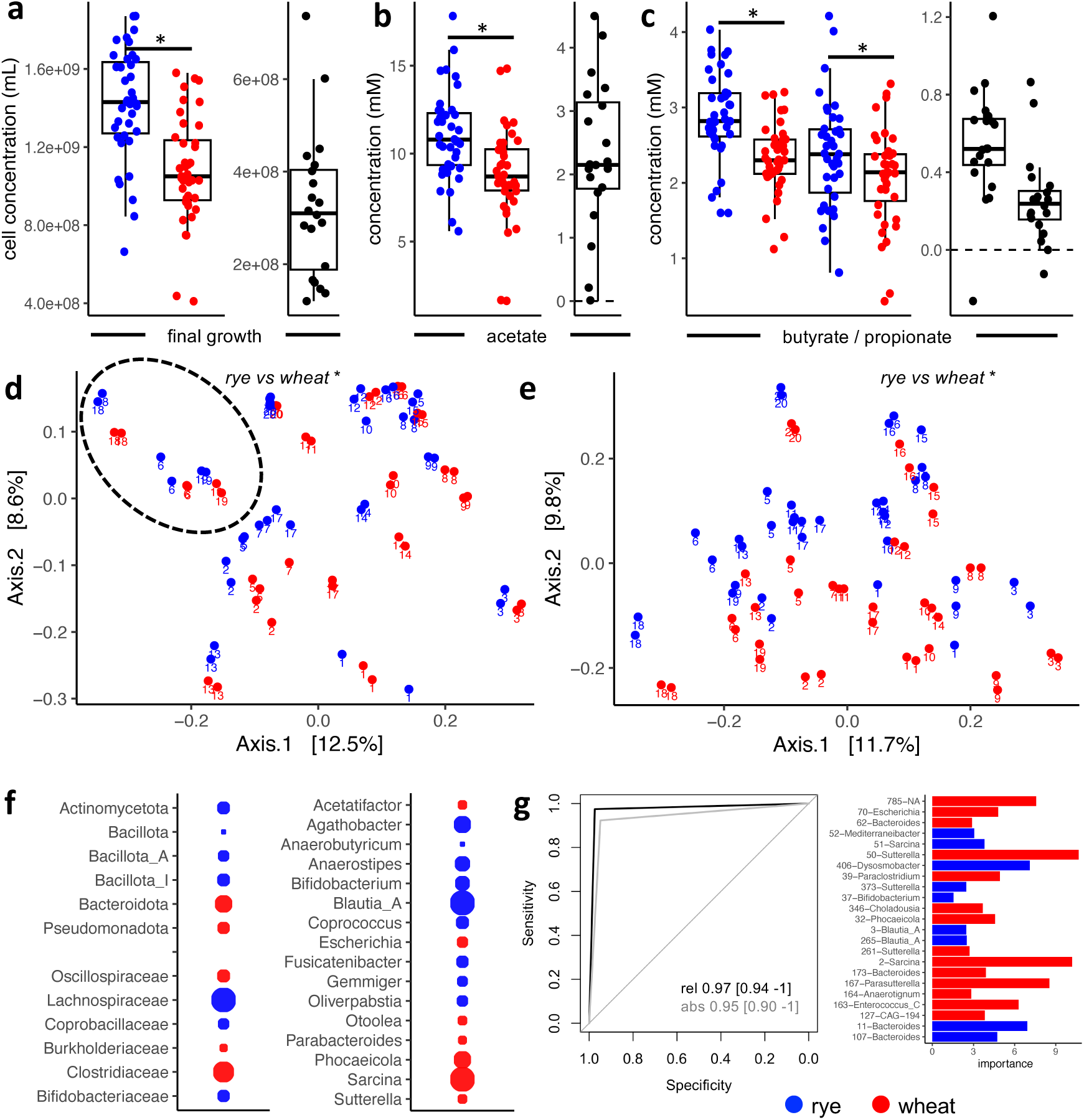
Growth of fecal communities derived from 20 individuals on pre-digested rye (blue) and wheat (red) for 24 hours; incubations were performed in replicate (n=2) samples. Panel a depicts final cell concentrations based on flow cytometric measurements, whereas panels b and c give acetate and butyrate / propionate concentrations formed, respectively; panels with black dots depict relative results (average values derived from rye subtracted by those derived from wheat). In panel d metric dimensional scaling analysis based on relative abundances on the ASV level (16S rRNA gene data) of grown communities is given with *Prevotella* dominated samples highlighted (n=3 individuals), whereas results based on absolute values are shown in panel e; IDs of individual subjects are displayed as well (communities of one individual (#4) were omitted due to their starkly contrasting composition) (*: p < 0.05 based on PERMANOVA results). Panel f depicts taxa significantly different between the two groups. Point sizes reflect estimates from linear regression analyses, where color depicts significantly (lfdr <0.05) higher final abundances in rye (blue) or wheat (red). Random forest analyses based on relative (rel) and absolute (abs) abundance data on the ASV level is given in panel g along with their AUC and 95% confidence intervals; importance values for ASVs included into the model based on relative abundance data is shown as well.

### Investigating bacterial growth dynamics on rye and wheat

We followed growth of fecal communities derived from five subjects over time (up to 63h) in order to investigate growth dynamics of gut bacteria on the two substrates. Ordination analyses based on 16S rRNA gene data demonstrated that community composition stayed subject-specific throughout the experiment. However, a significant influence of substrate based on PERMANOVA (p <0.05) was detected (Figure 2a). The composition of inocula clustered closely with the grown cultures of respective individuals. Four subjects displayed a *Bacteroides*-like enterotype, while one (P3) was dominated by *Prevotella*. Overall, bacterial multiplication was mainly observed during the first three time-points (till 23h), where final growth was reached in most samples (Figure 2b). For some samples slightly higher numbers were observed at 37h, whereas cell concentrations (minimally) declined after that time point. Differences in community composition were mostly established after 13h (t1) and were largely maintained throughout the experiment (Figure 2c). *Agathobacter*, *Anaerostipes*, *Anthropogastromicrobium*, *Blautia_A* and *Mediteraneibacter* were the most prominent genera enriched in rye, whereas *Bacteroides*, *Phocaeicola* and *Sarcina* were relatively higher abundant in wheat-derived samples; relative abundance of *Faecalibacterium* was only increased in rye until 18h (t2) (Figure 2c). The measured concentrations of butyrate were higher in rye than wheat, whereas no significant differences were detected for propionate in those experiments (Figure 2d). Calculated growth rates based on total cell concentrations were highest between 0h (t0) and 13h (t1) in cultures from both substrates (rye: 0.300 ± 0.026 h^-1^, wheat: 0.275 ± 0.0251 h^-1^) and strongly declined thereafter (t1>t2: 0.028 ± 0.018 h^-1^ (rye) and 0.032 ± 0.013 h^-1^ (wheat); t2>t3: 0.020 ± 0.015 h^-1^ (rye) and 0.023 ± 0.019 h^-1^ (wheat); t3>t4: 0.002 ± 0.005 h^-1^ (rye) and 0.008 ± 0.004 h^-1^ (wheat). Bacteria grew significantly faster (p<0.05) on rye compared with wheat in the first 13h and individual growth rates of most taxa were increased as well; only those of *Sarcina* were slightly higher in wheat containing cultures (Figure 2e). Correlation analyses of taxa based on their relative growth rate (growth rates derived from rye-growing cultures compared to those of wheat) clustered major butyrate producing taxa, such as *Agathobacter*, *Anaerostipes*, *Coprococcus* and *Faecalibacterium*, together along with *Blautia_A*, *Fusicatenibacter* and *Hominimerdicola* that were all negatively correlated with the wheat-enriched *Sarcina* (Figure 2f).

**Figure 2.**
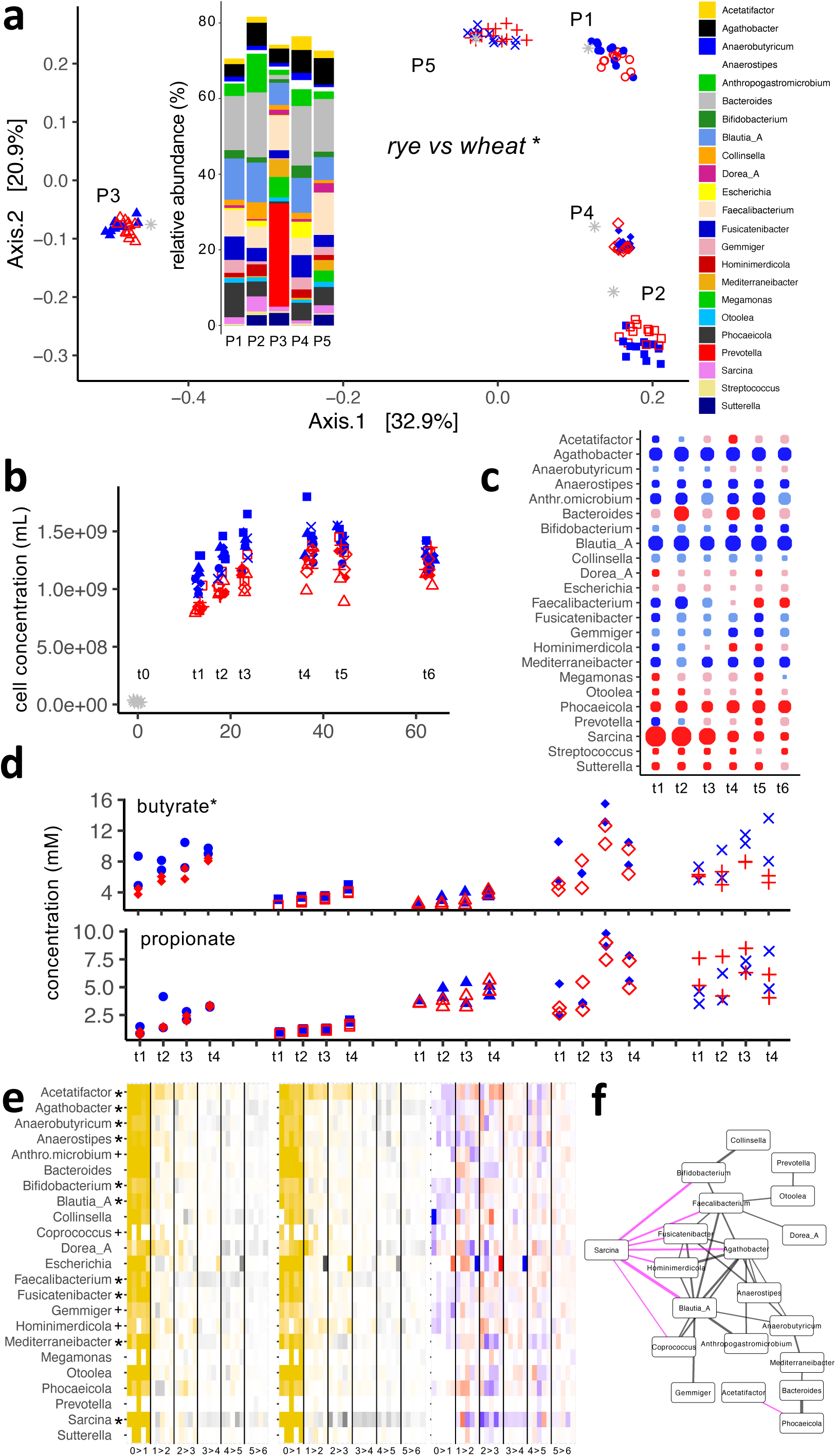
Monitoring dynamics of fecal communities derived from five individuals grown on pre-digested rye (blue) and wheat (red) for 63 hours; incubations were performed in replicate (n=2) samples. Panel a depicts metric dimensional scaling analysis of individual communities on the ASV level (16S rRNA gene data) at seven time-points including starting (inoculum) communities (shown as grey stars) (*: based on PERMANOVA results). Relative abundances of major genera is shown as well. Panel b depicts cell concentrations over time measured by flow cytometry, whereas relative abundance differences of major genera at individual time-points is given in panel c; dark colors depict significant (lfdr <0.05) results; point size reflects the estimate of linear regression analyses. In panel d measured concentrations of butyrate and propionate at four time-points is shown. In panel e calculated growth rates of major genera in rye (left) and wheat (middle), as well as relative growth rates (values of rye subtracted by those derived from wheat) (left) is given; for the first two panels gold and grey refer to positive and negative growth rates, respectively (*, +: lfdr < 0.05, < 0.1; growth rate differences between substrates in the first 13h). Panel f shows the correlation network based on relative growth rates including all values till 23h (t3) of incubation. Edge color and width depict correlation type (pink, negative; black, positive) and correlation strength, respectively.

### Metagenomics analyses of communities

Metagenomics analyses verified subject-specific clustering of samples, albeit significant differences between communities grown on the two substrates were detected (Figure 3a). In accordance, Bray-Curtis dissimilarities (BC) between subjects were higher than those between substrates within each individual at both time-points (t1 and t3). Increased relative abundances of the same taxa as observed based on 16S rRNA gene data, namely, *Agathobacter*, *Anaerostipes*, *Anthropogastromicrobium*, *Blautia_A* and *Mediteraneibacter,* along with *Wujia,* in rye and *Bacteroides*, *Phocaeicola* and *Sarcina* in wheat were observed*; Dorea, Eubacterium_F, Otoolea* and *Suterella* were additionally enriched during growth with the latter based on metagenomics data (Figure 3a, right panel; a detailed list of differentially abundant taxa is given in Table S2).

**Figure 3.**
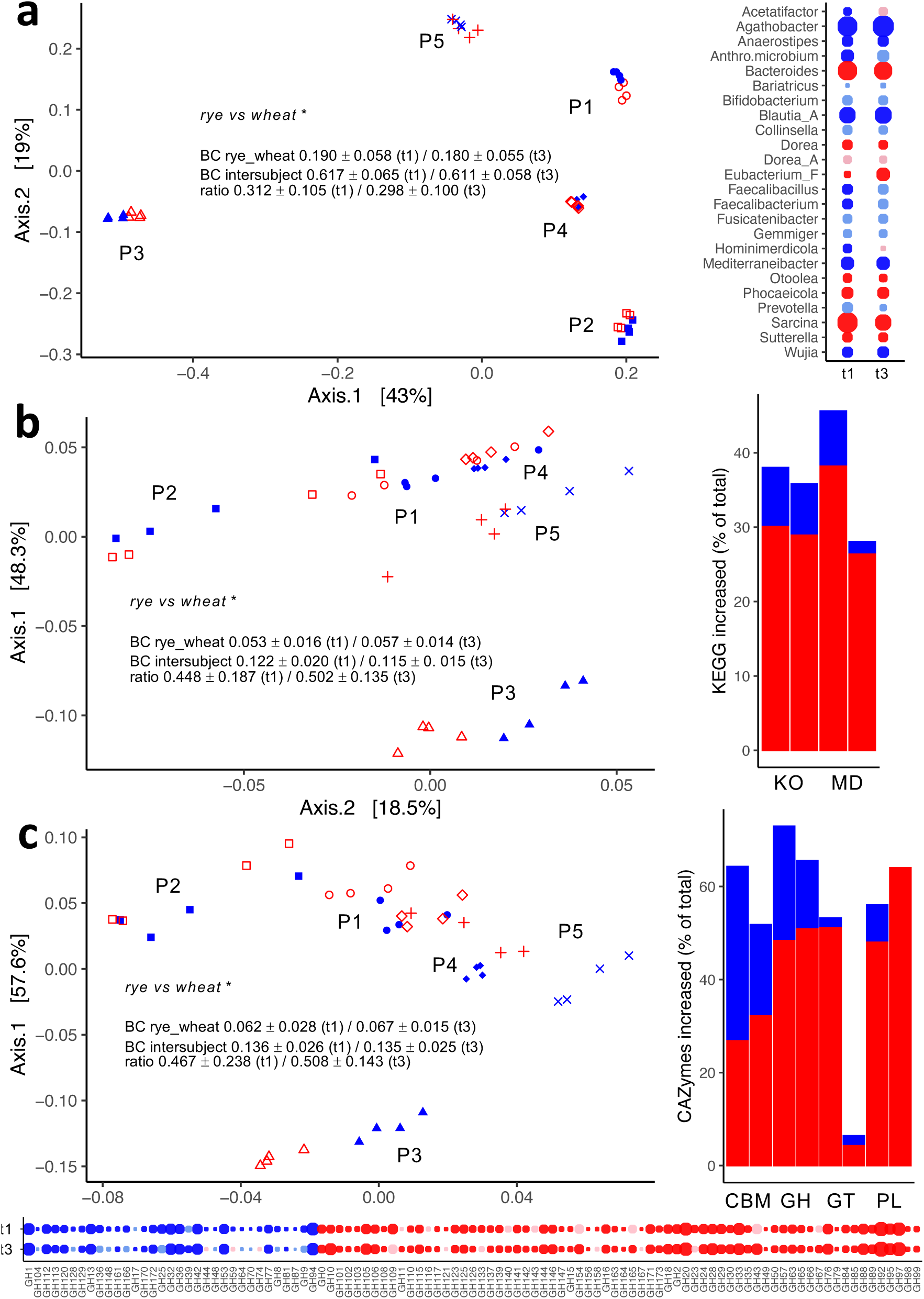
Metagenomic analyses of fecal communities derived from five individuals grown on pre-digested rye (blue) and wheat (red) for 13 hours (t1) and 23h (t3), respectively. Panel a depicts metric dimensional scaling analysis of individual communities on the species level; values of Bray Curtis dissimilarities (BC) between substrates (intra-subject) and between subjects as well as their ratios at both time-points is given as well. On the right, relative abundance differences of major genera at the two time-points is shown, where dark colors depict significant (lfdr <0.05) results; point size reflects the estimate of linear regression analyses. In panel b results based on KEGG orthologues (KO) are shown with percentage of significantly different (lfdr <0.05) KOs and KEGG modules (MD) given on the right (results from t1 are followed by those of t3). In panel c, respective results based on Carbohydrate Active Enzymes (CAZymes) is given (CBM: carbohydrate binding modules; GH: glycoside hydrolases; GT: glycoside transferases; PL: polysaccharide lyases). At the bottom, results from all GHs that were significantly different (lfdr <0.05) in relative abundance at any time-point are shown. *: p<0.05 based on PERMANOVA results.

Ordination analyses based on KEGG orthologues (KO) also revealed strong subject-specific signatures; similar to taxonomic analysis above samples from the *Prevotella*-enriched samples clustered most separately from others (Figure 3b). The ratio of BC between intra- and inter-subject comparisons was, however, significantly (p <0.05) higher compared to that based on taxonomic data indicating that, on a functional level, the influence of substrate on community structure was increased. A multitude of KOs and KEGG modules (MD) were significantly differently abundant between samples derived from the two substrates with many functions higher in the wheat group (Figure 3b, right panel; a detailed list of differentially abundant KOs and MDs is given in Table S3).

Analyses based on Carbohydrate Active Enzymes (CAZYmes) showed a similar picture as those based on KOs. Subject-specific signatures dominated, however, the BC ratio was significantly increased compared to taxonomy-based results (Figure 3c). A high percentage of CAZymes of individual families was significantly differentially abundant between substrate groups with many being enriched in wheat-derived samples (Figure 3c, right panel; a detailed list of differentially abundant CAZymes is given in Table S4). Detailed analyses of Glycoside Hydrolases (GH), which are key enzymes for the degradation of (complex) carbohydrates, revealed strong differences between groups with 30 (t1) / 60 (t3) and 18 (t1) / 62 (t3) significantly enriched in rye and wheat, respectively, representing 74.4% (t1) / 65.6% (t3) of all detected GHs (Figure 3c, bottom panel). Results demonstrate distinctly different carbohydrate degradation machineries between communities grown on the two substrates.

### Metatranscriptomics analyses of communities

In order to obtain insights into the gene-expression level we performed metatranscriptomics analyses of actively growing communities at t1 (13h). Based on taxonomic results subject-specific patterns also dominated on this level (Figure 4a). Number of taxa that were significantly different between the two substrate groups was lower compared to DNA-based analyses. For major genera (species) only *Agathobacter* (main members were *A. faecis* and *A. rectale*) and *Agathobaculum* (main species *A. butyriproducens*) were significantly enriched in rye and wheat, respectively (Figure 4a, right panels; a detailed list of differentially abundant taxa is given in Table S5).

**Figure 4.**
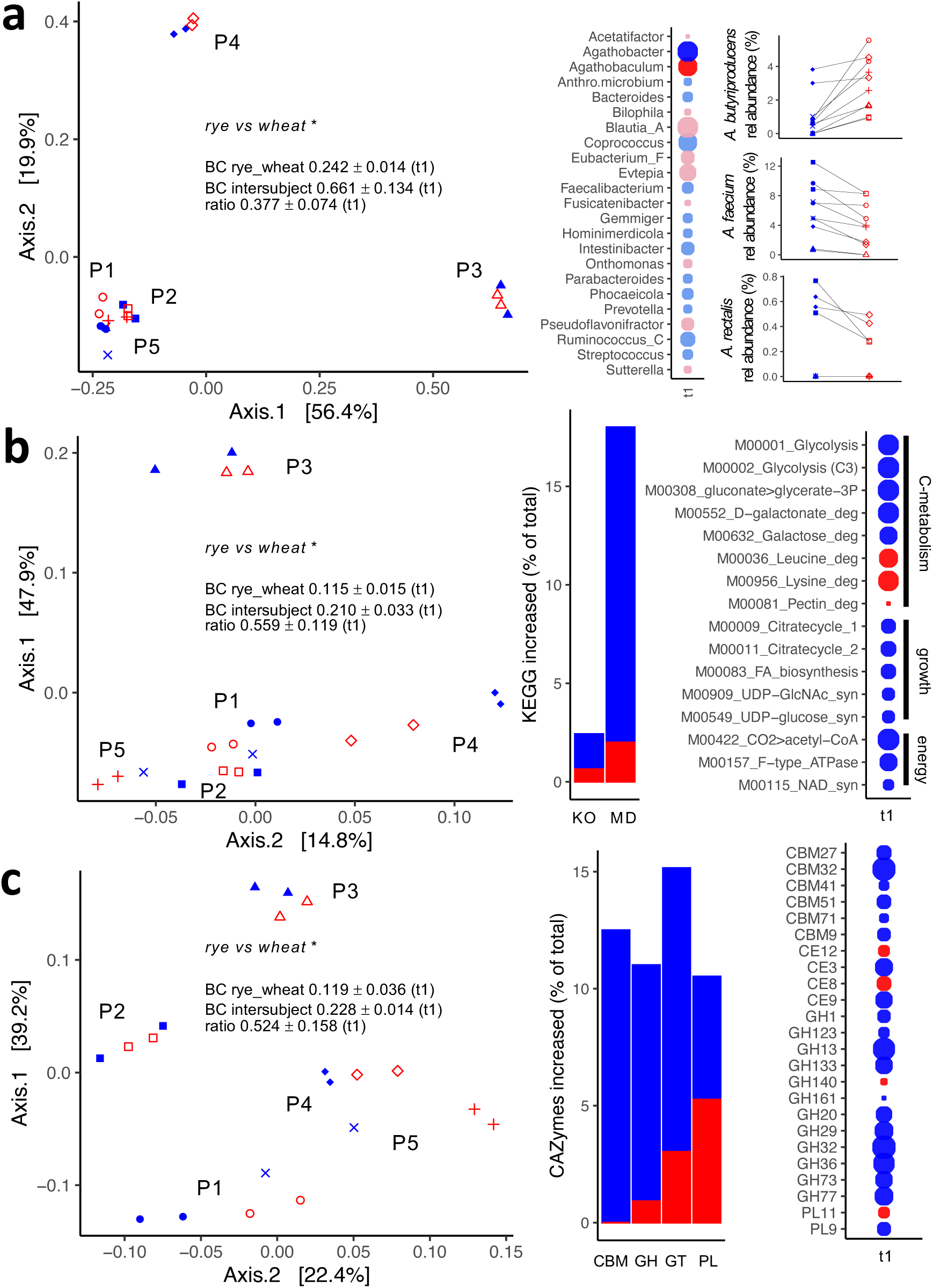
Metatranscriptomics analyses of fecal communities derived from five individuals growing on pre-digested rye (blue) and wheat (red) for 13 hours (t1). Panel a depicts metric dimensional scaling analysis of individual communities on the species level; values of Bray Curtis dissimilarities (BC) between substrates (intra-subject) and between subjects as well as their ratios is given as well. On the right relative abundance differences of major genera is shown, where dark colors depict significant (lfdr <0.05) results; point size reflects the estimate of linear regression analyses. On the very right individual relative abundance data of *Agathobaculum butyriproduces*, *Agathobacter faecis* and *Agathobacter rectalis* is shown. In panel b results based on KEGG orthologues (KO) are shown with percentage of significantly different (lfdr <0.05) KOs and KEGG modules (MD) given on the right; results of major significantly different MDs are shown on the very right. In panel c, respective results based on Carbohydrate Active Enzymes (CAZymes) is given (CBM: carbohydrate binding modules; GH: glycoside hydrolases; GT: glycoside transferases; PL: polysaccharide lyases); on the very right results of individual CAZymes significantly different (lfdr <0.05) between the two substrate groups are shown. *: p <0.05 based on PERMANOVA results.

On the functional level based on KOs, samples still predominantly clustered based on subjects (Figure 4b). However, the BC ratio was significantly higher compared to taxonomy results based on gene expression, as well as to those based on KOs on the DNA level (Figure 3b), suggesting that the individual substrate strongly influenced gene-expression of communities on a functional level. Expression of several KOs and MDs were significantly different between samples derived from the two groups with most enriched in the rye group (Figure 4b, right panel; a detailed list of differentially abundant KOs and MDs is given in Table S6). More detailed analyses demonstrated that communities growing with rye expressed genes associated with growth and energy generation as well as carbohydrate degradation at higher levels indicating more active communities compared to those growing with wheat; expression of genes linked to degradation of certain amino acids and pectin was increased in the wheat group (Figure 4b, most right panel).

On the CAZyme level similar patterns in ordination analysis as found with KOs were observed (Figure 4c) with many CAZymes higher expressed in the rye group (Figure 4c, right panel). In the case of GHs, they were almost explicitly more expressed in communities growing on rye supporting strong carbohydrate degradation activities in those communities (Figure 4c, most right panel). Among them were classical α-amylases (GH13 and GH77) that degrade (resistant) starch-like compounds, a α-fucosidase (GH29) and GH32, which was the most differentially expressed GH and acts on fructose containing polysaccharides including fructans.

### Detailed analyses of SCFA-producing pathways

A major focus of this study was to investigate the production of SCFA and we, hence, took a detailed look into respective pathways using metagenomics and metatranscriptomics data. While acetate is produced by most members of gut microbiota only certain microbes are forming butyrate and propionate, respectively. Based on relative abundance data genes of the main butyrate-synthesis pathway, the so-called acetyl-coA pathway (accoa), and the gene encoding the pathways’ most important terminal enzyme, i.e., butyrate transferase (*but*), were not differentially abundant between substrate groups (Figure 5a,b). At both time-points the gene encoding butyrate kinase (*buk*) and the main propionate-producing pathway, which is characterized by the intermediate compound succinate (suc), were increased with wheat, whereas the propandiol (pdiol) pathway that yields propionate as well was relatively higher abundant in communities growing with rye (Figure 5a,b). The relative abundance of accoa and *but* increased from t1 to t3 in both substrate groups; no differences between time-points were detected for other pathways (Figure 5b).

**Figure 5.**
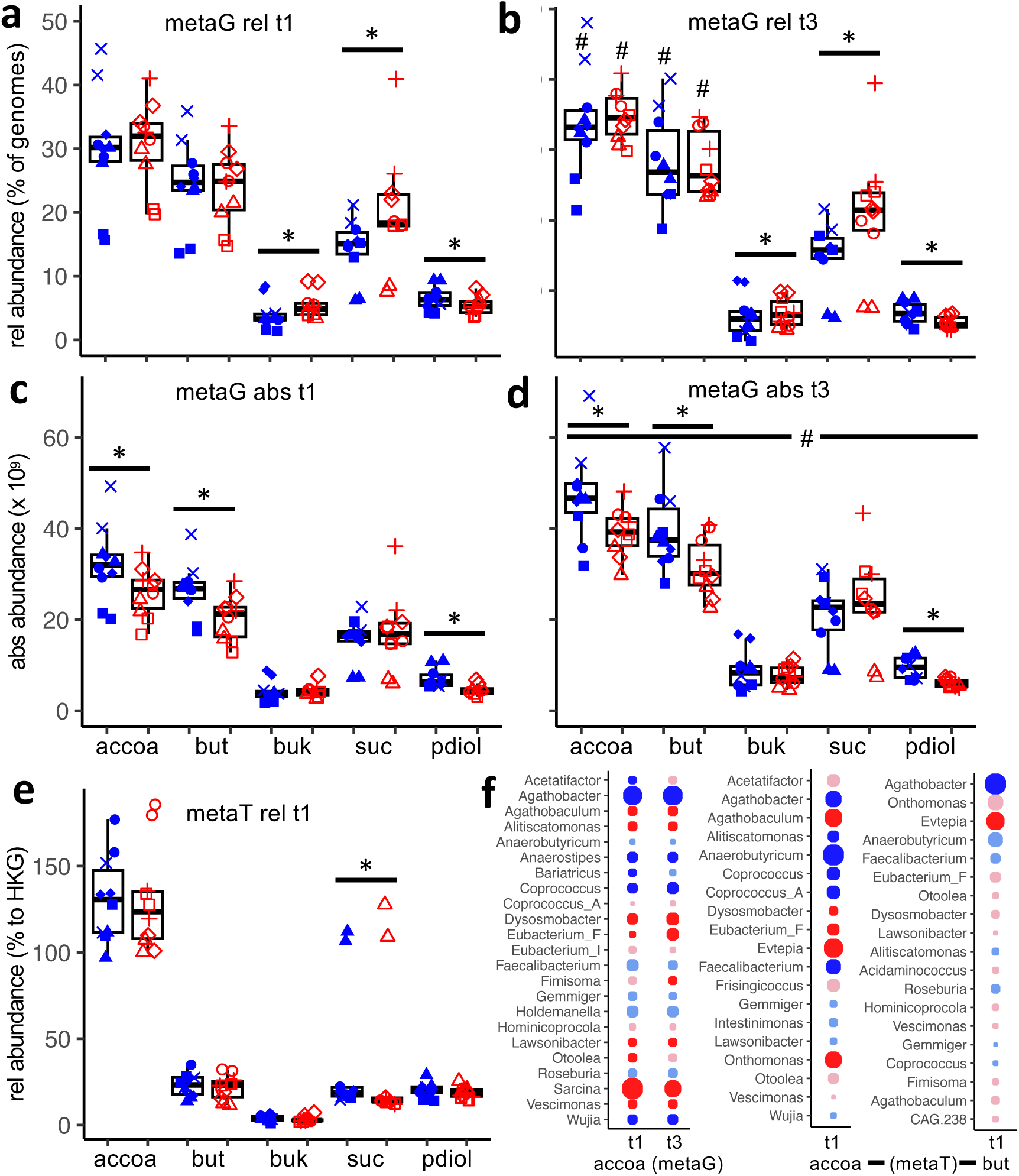
Detailed analysis on butyrate and propionate synthesis pathways based on metagenomics (metaG) and metatranscriptomics (metaT) data of fecal communities derived from five individuals (replicate samples; n=2) grown on pre-digested rye (blue) and wheat (red). Abundance of genomes exhibiting pathways relative to all genomes after 13 hours (t1) and 23 hours (t3) of growth is shown in panels a and b respectively, whereas respective absolute abundance data is given in panels c and d. Results of the main butyrate synthesis pathway (accoa) along with its two major terminal enzymes, butyrate transferase (but) and butyrate kinase (buk), as well as those of the two dominant propionate producing pathways with succinate (suc) and propandiol (pdiol) as intermediate signature compounds are shown (*, #: significantly different between groups, time-points). In panel e, metatranscriptomics results of individual pathways relative to average expression of housekeeping genes (HKG) is given. In panel f results of taxa comprising the butyrate pathway are displayed, where point size reflects the estimate of linear regression analyses and dark colors depict significant (lfdr <0.05) differences; for metatranscriptomics data results for the entire accoa and for the terminal enzyme butyrate transferase (but) are shown separately.

Based on absolute abundance data the accoa and *but*, along with pdiol, were significantly higher abundant in communities growing with rye at both time-points (Figure 5c,d). Due to overall growth of bacteria, all pathways were higher at t3 compared with t1 (Figure 5d). On the gene-expression level no difference for any pathway was detected between substrate groups, except for suc that was higher expressed in the rye group (Figure 5e). Overall, genes associated with accoa were much higher expressed (relative to house- keeping genes) than those encoding for terminal enzymes and genes comprising propionate pathways (Figure 5e).

Detailed analyses of butyrate producers suggested *Agathobacter* as the main (differentially abundant) taxon in the rye group, along with *Anaerostipes*, *Coprococcus* and *Wujia*, whereas *Sarcina,* together with *Agathobaculum*, *Dysmosobacter* and others, were relatively higher abundant in the wheat group (Figure 5f, left). Based on expression data of the accoa pathway *Anaerobutyricum*, *Coprococcus, Coprococcus_A* and *Faecalibacterium* were also increased in the rye group; *Evtepia* and *Onthomonas* were revealed to be specific for the wheat group (Figure 5f, middle panel). Based on expression of the key gene *but* only *Agathobacter* (higher in rye) and *Evtepia* (higher in wheat) were significantly different between substrate groups (Figure 5f, right panel).

### Gene expression patterns of Agathobacter faecis based on metagenome assembled genomes

*Agatobacter faecis* was revealed as the key taxon for higher butyrate production in communities growing with rye in above analyses and we, hence, took a more detailed look into its physiology mapping gene-expression data on metagenome assembled genomes (MAGs) of this species. For all individuals one high-quality MAG (completeness: 97.7 ± 1.6%; contamination 2.0 ± 0.67%) that was annotated as *A. faecis* was obtained. Ordination analysis based on gene-expression data including all genes shared between all genomes (n=1.773) displayed clear subject-specific signatures with those from P3 and P5 clustering distant to other samples (Figure 6a). Phylogenetic relatedness can only partly explain this observation; the genome from P3 was distantly related to others, however, that from P5 showed high ANI similarities to others despite its unique expression pattern (Figure 6a, dendrogram). Overall, expression patterns were significantly influenced by substrate and several genes as well as functions based on KOs, MDs and CAZymes were differently expressed between the two groups (p <0.05; results were based on raw p-values only as none were detected significantly different after multiple testing correction). Detailed analyses revealed two key genes that were considered important for rye degradation higher expressed in the rye group in all individuals (Figure 6b). The first is likely involved in arabinoxylan degradation as it encodes a carbohydrate binding module showing affinity for xylans (CBM91) fused to a sequence encoding an α-L-arabinofuranosidase (GH43) (Figure 6b, upper panel). The other gene encodes the fructose phosphotransferase system that catalyzes the uptake of fructan-derived monomers (Figure 6b, lower panel). Investigating gene expression of individual genes comprising the accoa pathway as well as those encoding the enzymes for acetate formation, namely, phosphotransacetylase (*pta*) and acetate kinase (*ackA*) did not reveal any clear distinctions between substrate groups suggesting that pattern of SCFA formation was not influenced by the substrate *per se* for this taxon (Figure 6c).

**Figure 6.**
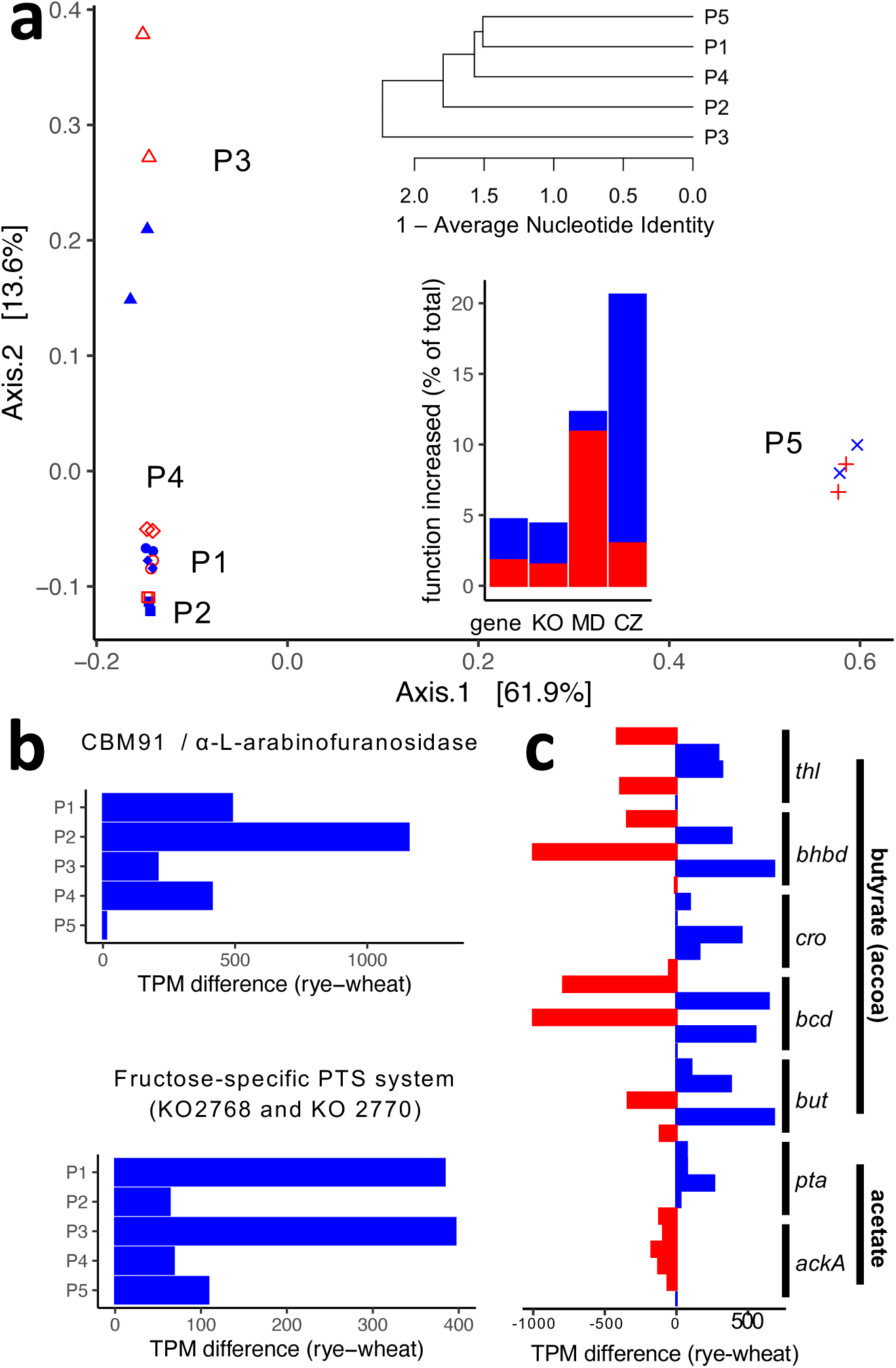
Detailed analyses of *Agathobacter faecis* genomes assembled from samples of individual subjects. For each subject one high quality metagenome assembled genome (MAG) was reconstructed and their gene expression patterns at t1 (13 hours of growth) based on metatranscriptomics data were investigated. Panel a depicts metric dimensional scaling analysis of expression data based on genes shared by all five genomes (n=1.733). Phylogenetic relationships of genomes based on the Average Nucleotide Identity (ANI) along with percentages of significantly (p <0.05) differentially expressed genes, KEGG orthologues (KO) and modules (MD) as well as CAZymes (CZ) between groups are shown as well. Panel b depicts detailed expression results based on genes encoding (i) an enzyme involved in arabinoxylan degradation (upper panel) and (ii) the fructose PTS uptake system (lower panel); relative data (values form rye subtracted by those from wheat) based on average values of replicate samples (n=2) of individual subjects are shown. In panel c, relative expression data for all genes of the butyrate (thl: thiolase, bhbd: β-hydroxybutyryl-CoA dehydrogenase, cro: crotonase, bcd: butyryl-CoA dehydrogenase, but: butyryl-CoA:acetate CoA-transferase) and acetate (pta: phosphotransacetylase, ackA: acetate kinase) synthesis pathways of individual subjects (P1 to P5) are given.

## Discussion

With this study we were able to uncover distinct microbial features of human gut microbiota involved in degradation of wheat and rye. A strength of our study were the multi-level analyses involving quantitative insights into community composition, their growth dynamics as well as details on functions comprising both the DNA and RNA level. Throughout experiments person-specific signatures were prevailing, which was in particular true for compositional data. However, despite individual microbial backgrounds, common responses to each substrate were observed, especially on a functional level. The results demonstrate that functional redundancy, i.e., the same function is encoded on various taxonomies, plays an important role to adapt to (varying) nutritional conditions despite differences in taxonomic make-ups ^22,23^. This was exemplified by glycoside hydrolases, where over 65% of all families detected were differentially abundant between communities grown on rye and wheat, respectively. This demonstrated strong differences in their carbohydrate degrading enzyme spectra. Data on gene expression showed the strongest substrate-specific signatures, where BC ratios between inter-subject communities and those comparing results within subjects were the lowest. For instance, several functions involved in substrate degradation, particularly enriched in rye, such as GH32 and CE3 acting on arabinoxylan, were detected highlighting the value of expression data to unravel microbial features specific for certain nutritional conditions.

Overall, our *in vitro* data are in accordance with corresponding *in vivo* observations, where the plasma SCFA acetate and butyrate were increased upon rye interventions, along with the butyrate producer *Agathobacter* ^21,24^, demonstrating that *in vitro* models are useful to investigate degradation mechanisms of substrates by gut microbiota and that results can be extrapolated to *in vivo* settings. This is also true for other fibers. For instance, we could recently reveal that *Anaerostipes* and *Faecalibacterium* are the main drivers for an increased butyrate production potential upon inulin administration in dietary intervention studies, whereas *Agathobacter* is the butyrate producer most stimulated by resistant starches ^9^. Those signature taxa for individual substrates were also revealed *in vitro* earlier ^4^. It underscores the potential of *in vitro* methods for screening of a multitude of substrates using the same inoculum, which is hardly feasible in real-world intervention studies. Furthermore, *in vitro* experiments allow to investigate the degradation dynamics of individual taxa and discern associated functions on multiple levels in detail as done here. In this context, our metagenomics/metatranscriptomics approach allowed annotation of assembled full-length genes on the DNA level that formed the basis for robust analyses of gene expression data. Furthermore, it enabled genome-resolved insights, as exemplified by analyses of the key butyrate producer *Agathobacter faecis (*former *Roseburia faecis)* providing detailed insights into its physiology during growth in a complex community. Overall, neither on the DNA nor the RNA level the abundance of the main butyrate-forming pathway was relatively increased in the rye group. It was, however, increased in absolute terms suggesting that higher production of this SCFA is primarily due to better substrate availability of rye that promoted faster and higher overall growth of bacteria. Also in the case of *A. faecis* clear adaptations to rye degradation were discovered, however, we did not find any physiologic shifts regarding SCFA formation compared with wheat. We additionally found propionate increased in the rye group, along with the pdiol pathway, which was not observed *in vivo* ^21^. SCFA concentrations measured in plasma are often distinct to those in the intestinal environment due to their consumption at various body sites and concentrations measured in the circulation do, hence, not reflect their true production quantities ^25^. While butyrate is used by the colonic epithelium for energy generation through β-oxidation, relatively more propionate reaches the liver and can be used for gluconeogenesis ^26^. This can be particularly important during weight-loss interventions, as performed in the mentioned *in vivo* reference study above, and might explain why propionate was not found at increased levels in the circulation.

*In vivo*, the upper gastrointestinal tract acts on ingested substrates before they reach the colon and pre-digestion is, hence, important in order to study the degradation of substrates by (colonic) gut microbiota *in vitro*. In the case of fibers this can often be neglected due to their nature being resistant to digestion in upper parts of the intestines. However, for whole foods, including wheat and rye, only a fraction is reaching the colon with strong modifications of their nutrient composition (Table 1). We showed previously that incubating fecal communities of pigs with raw and pre-digested substrates, respectively, resulted in stark compositional differences in grown-up communities (*preprint coming soon*). Our approach to pre-digest substrates in Minipigs can be regarded as a strength of this study as pigs’ digestive physiology resembles closely that of humans making them good model animals for nutrition research ^27,28^. However, it also came with downsides as some indigenous bacteria from pigs were introduced together with the pre-digested substrates into experimental cultures, despite lyophilization and disinfection with ethanol before use. This was in particular true for *Sarcina* as this taxon was not found in fecal inocula or in cultures grown on baseline medium. Results associated with this taxon should, hence, be interpreted with care, however, we do not think that it interfered with main results found in this study.

Several studies have shown amelioration of cardiometabolic parameters upon rye ingestion ^29,30^ and our study substantiates beneficial host effects of this cereal mediated by gut microbiota via increased SCFA production. The results obtained were enterotype independent as *Prevotella* enriched cultures showed a similar increase in cell and SCFA concentrations, along with higher abundances of *Agathobacter,* in rye cultures compared to those containing wheat. For individual fibers this is often not the case, where community structure can strongly influence outcomes ^20,31^. The complex substrate nature of rye containing distinct carbohydrates might balance out differences and increase its applicability to promote SCFA production in broad groups of the population.

Next to increased qualitative characteristics of rye in terms of SCFA synthesis its precaecal digestibility of the carbohydrate fraction is also lower compared to that of wheat, where, overall, more substrate reaches the large intestine and becomes available to gut microbiota ^18^. The increased filling of the colon and the ongoing fermentation processes prolong the feeling of satiety due to the so-called “colonic brake,” where satiety hormones such as peptide YY are released, slowing down further gastric emptying ^32^. As a result, a preventive effect against obesity is postulated and a diet with a high share of rye indeed led to greater weight loss than a diet with a high share of wheat ^33^. Furthermore, increased amounts of substrates reaching the colon probably further enhance the production of SCFA from rye compared to wheat.

In summary, our study provides crucial knowledge on functions of gut microbiota involved in the degradation of rye and wheat, two leading bread cereals, at multiple levels and uncovered molecular mechanisms underlying increased SCFA production capacity of the former. It substantiates that the observed health benefits of rye in dietary intervention studies are also mediated by gut microbiota.

## Materials and Methods

### Cultivation of fecal communities

Cultivation was done similar as described previously ^4^ with modification. In brief, fresh stool samples were diluted to yield ∼1×10^7^ cell mL^-1^ starting concentrations in cultures harboring anaerobic YcFA medium (pH = 6.8) in Hungate tubes (V=10 mL) containing 2 g L^-1^ of substrate rye or wheat, respectively. The medium is described in reference and is based on YCFA, however, with reduced amounts of yeast extract and casitone (0.5 g L^-1^ each) in order to minimize background carbon sources. Pre-digestion of substrates was achieved as described earlier (*preprint coming soon*) by feeding ileo-caecal fistulated Göttingen minipigs with 100% of ground rye or wheat, respectively, and ileal digesta were collected (each pigs received both cereals with an in-between break of at least one week). The digesta of three pigs was pooled and lyophilized. Before use, the substrate was treated with ethanol (100%) for 10 minutes in order to limit growth of accompanying bacteria. Cultures were incubated for 24h (rpm 200) at 37 °C. For time-series and metaOmics experiments the same set-up was upscaled to V=100 mL using Schott bottles with caps providing anaerobic conditions. Samples were taken at 13h (t1), 18h (t2), 23h (t3), 37h (t4), 44h (t5) and 63h (t6).

### Sampling and measurement of cell concentration and SCFA

Samples were taken to measure cell concentrations by flow cytometry and 16S rRNA gene analysis as described previously ^4^. Furthermore, an aliquot was subjected to SCFA analyses by GC-MS as described previously ^34^ SCFA measurement of samples from Schott flasks (time-series experiments) was done according to Kircher and colleagues ^4^. However, values for acetate were not usable due to a problem with the internal standard. For metagenomics (t1 and t3) the same DNA as for 16S rRNA gene analyses was used. For metatranscriptomics (t1) 1 mL of sample was immediately added to 1 mL RNA later (Thermo Fisher Scientific, USA) centrifuged and resuspended in 0.5 mL RNA later before storage at -80 °C.

### Nucleic acid extraction, library preparation and sequencing

DNA was extracted using the ZymoBIOMICS Miniprep Kit (ZYMO, USA) according to the manufacturer. Amplification of the V3V4 region of the 16S gene and subsequent sequencing on Illumina MiSeq (2x 250 bp) was done as before ^35^. For metagenomics, libraries were prepared (Illumina DNA Prep, Illumina, USA) and products were subsequently sequenced on NovaSeq 6000 (2x 150 bp) as described previously ^35^. For metatranscriptomics, RNA was extracted (RNeasy Mini Kit, Qiagen, Germany) followed by DNase I treatment (RNase-Free DNase Set, Qiagen, Germany). Depletion of ribosomal reads and subsequent library preparation was done according to the manufacturer (Ribo-Zero Plus, Illumina, USA) followed by NovaSeq 6000 sequencing.

### Bioinformatics analyses

For 16S rRNA gene analysis raw sequences were analysed in R (v4.2.2) using DADA2 (v1.20) ^36^ as described previously ^4^. Amplicon sequence variants (ASVs) were annotated based on GTDB (r2.20). For metagenomics, a genome-resolved workflow based on MetaWRAP (v1.3.0; default mode) ^37^ was applied as described in Kircher and colleagues ^4^ based on MEGAHIT (assembly) and MaxBin 2, MetaBAT 2 and CONCOCT (binning); all samples derived from each individual were assembled together and later separated in the binning process. For taxonomic analyses (based on GTDB r2.20) reads were mapped via bwa-mem (0.7.17-r1188; default mode) ^38^ to house-keeping genes (HKG; based on gtdb-tk, v2.1.0) ^39^ derived from high-quality (completeness >80%; contamination <5%) metagenome assembled genomes (MAGs) together with those of all species of the Unified Human Gastrointestinal Genome (UHGG.v2; only species represented by an isolate or more than three MAGs were considered) ^40^. Functional annotation of proteins was done via prokka (v1.14.6; --fast --norrna --notrna – metagenome) ^41^ on output of assembled contigs and were based on KEGG Orthologs (KO) and Carbohydrate Active Enzymes (CAZymes) using KofamScan (v1.3.0; default mode) ^42^ and dbCAN4 (v2.0.11; default mode) ^43^, respectively. Reads were mapped to all contigs using bwa-mem filtered by msamtools (v1.1.3; filter -b -l 70 -p 95 -z 80) and subsequently featureCounts (v2.0.3; default mode) ^44^ was used to call number of reads mapped to individual genes. Raw reads were first mapped onto assembled contigs and unaligned reads were subsequently mapped against genomes of representative species of the UHGG.v2. Results were merged, gene-length corrected (RPKs) and normalized against average results of HKGs yielding genome normalized results as described previously ^4^. Specific analyses of SCFA pathways was done as described earlier ^4^, where reads were mapped against a catalogue of pathway genes derived from MAGs and UHGG.v2 representatives using bwa-mem; results were gene-length corrected and normalized to HKGs yielding genome normalized results as above.

For metatranscriptomics analyses, reads were quality filter and depleted of ribosomal reads using Kneaddata (Huttenhower lab; v0.7.2). The same workflow as described above for metagenomics was then applied, where taxonomic results were based on HKGs, whereas functional insights were obtained mapping reads against annotated gene sequences (KO, CAZymes and SCFA pathways). As for metagenomics analyses, results were gene-length corrected and expressed relative to average results of HKGs. For specific analyses of *Agathobacter faecis* reads were mapped to genes of respective MAGs (one per individual was obtained) using bwa-mem and subsequently filtered by msamtools. The Average Nucleotide Identity (ANI) of the five MAGs was calculated based on fastani (v1.32; default mode) ^45^. They were then subjected to pangenome analysis using roary (v3.13.0; default mode) ^46^ and results, expressed as transcripts per million (TPMs) ^8^, of only genes shared by all genomes (n=1.773) were considered in subsequent analyses.

### Data analysis

Statistical analyses were performed in R (v4.2.2) applying linear (mixed-effect) models (function *lmer* from the *lme4* (v1.1-34) package) using log transformations (log (data+1)) and including subject as a random effect. All taxa with an average abundance of 1% (for species the cut-off was lowered to 0.1%) were included into analyses, whereas for functional features a cut-off of >50% presence in all samples was applied. All results were corrected for multiple testing using *fdrtools* (v1.2.17), where a lfdr <0.05 was considered significant unless explicitly stated otherwise; for transcriptomics results of *A. faecis* MAGs a raw p-value <0.05 was considered. Ordination analyses are based on metric multidimensional scaling (MDS) using the *phyloseq* package (v1.36.0) ^47^. Bray Curtis dissimilarities (BC) were calculated using the function *vegdist* from the *vegan* package (v2.5.7; doi: 10.32614/CRAN.package.vegan), whereas PERMANOVA analyses were performed via the function the adonis2 from the same package. For absolute abundance data proportional count data were multiplied by bacterial concentrations and subsequently analyzed the same way as above for relative abundance data. Growth rates (μ) were calculated from absolute abundance data of individual taxa on the genus level (μ = (ln abundance t2 - ln abundance t1) / (t2 – t1)). All plots were constructed in R via ggplot2 (v3.3.5). Random Forest analyses were done using the *caret* package (v6.0-94; method=”ranger”) ^48^ applying a five-fold cross split, five-fold repeat strategy to avoid overfitting. Features were pre-selected based on abundance (presence in 20% of samples) and *boruta* (v8.0) analysis ^49^. ROC-curves and AUC calculations were done based on the package *pROC* (v1.18.4) ^50^.

## Funding

The project was supported by funds of the Federal Ministry of Food and Agriculture (BMEL) based on a decision of the Parliament of the Federal Republic of Germany via the Federal Office for Agriculture and Food (BLE) under the innovation support program (281B101016). This work was also funded by the DFG (project #456214861). Marius Vital was additionally funded by HiChol (01GM2204).

## Ethics approval

The animal trials were conducted in accordance with German regulations and approved by the Ethics Committee of Lower Saxony for the Care and Use of Laboratory Animals (LAVES: Niedersächsisches Landesamt für Verbraucherschutz und Lebensmittelsicherheit; reference: 33.8-42502-04-18/2884). The use of fecal samples was approved by local ethic authorities (#8566_BO_K_2019) and all subjects have given informed consent.

## Disclosure Statement

The authors report no conflict of interest.

## Data Availability Statement

All sequences are publicly available at the European Nucleotide Archive (*PRJ coming soon*).

## Authors’ contributions

M.V., C.B.H., C.V., A.v.F. and A.K. conceived the study. C.B.H., A. K., K.E.B.K., S.W., R.G. performed laboratory analyses. M.V. performed bioinformatics analyses. M.V., C.B.H. and A.K. performed data analysis and interpreted the results. M.V. and C. B. H. wrote the manuscript. All authors reviewed and edited the manuscript.

## Acknowledgements

We want to thank all participants involved in the study for providing fresh sample.

## Supplemental online material

**Table S1.** Differentially abundant taxa after growth on the two substrates based on 16S rRNA gene data. The estimated effect size (Estimate) from linear regression analyses including subject as a random effect is shown, where negative and positive values refer to higher abundance in rye and wheat, respectively. Raw p-values along with the lower false discovery rate (lfdr) after multiple testing correction is shown.

**Table S2.** Differentially abundant taxa after growth on the two substrates (t1 and t3 combined) based on metagenomics data. The estimated effect size (Estimate) from linear regression analyses including subject as a random effect is shown, where negative and positive values refer to higher abundance in rye and wheat, respectively. Raw p-values along with the lower false discovery rate (lfdr) after multiple testing correction is shown.

**Table S3.** Differentially abundant KEGG Orthologs (KO) and modules (MD) after growth on the two substrates (t1 and t3 combined) based on metagenomics data. The estimated effect size (Estimate) from linear regression analyses including subject as a random effect is shown, where negative and positive values refer to higher abundance in rye and wheat, respectively. Raw p-values along with the lower false discovery rate (lfdr) after multiple testing correction is shown.

**Table S4.** Differentially abundant CAZYmes after growth on the two substrates (t1 and t3 combined) based on metagenomics data. The estimated effect size (Estimate) from linear regression analyses including subject as a random effect is shown, where negative and positive values refer to higher abundance in rye and wheat, respectively. Raw p-values along with the lower false discovery rate (lfdr) after multiple testing correction is shown.

**Table S5.** Differentially abundant taxa after growth on the two substrates based on metatranscriptomics data. The estimated effect size (Estimate) from linear regression analyses including subject as a random effect is shown, where negative and positive values refer to higher abundance in rye and wheat, respectively. Raw p-values along with the lower false discovery rate (lfdr) after multiple testing correction is shown.

**Table S6.** Differentially abundant KEGG Orthologs (KO) and modules (MD) after growth on the two substrates based on metatranscriptomics data. The estimated effect size (Estimate) from linear regression analyses including subject as a random effect is shown, where negative and positive values refer to higher abundance in rye and wheat, respectively. Raw p-values along with the lower false discovery rate (lfdr) after multiple testing correction is shown.

**Table S7.** Differentially abundant CAZYmes after growth on the two substrates based on metatranscriptomics data. The estimated effect size (Estimate) from linear regression analyses including subject as a random effect is shown, where negative and positive values refer to higher abundance in rye and wheat, respectively. Raw p-values along with the lower false discovery rate (lfdr) after multiple testing correction is shown.

**Figure S1.**
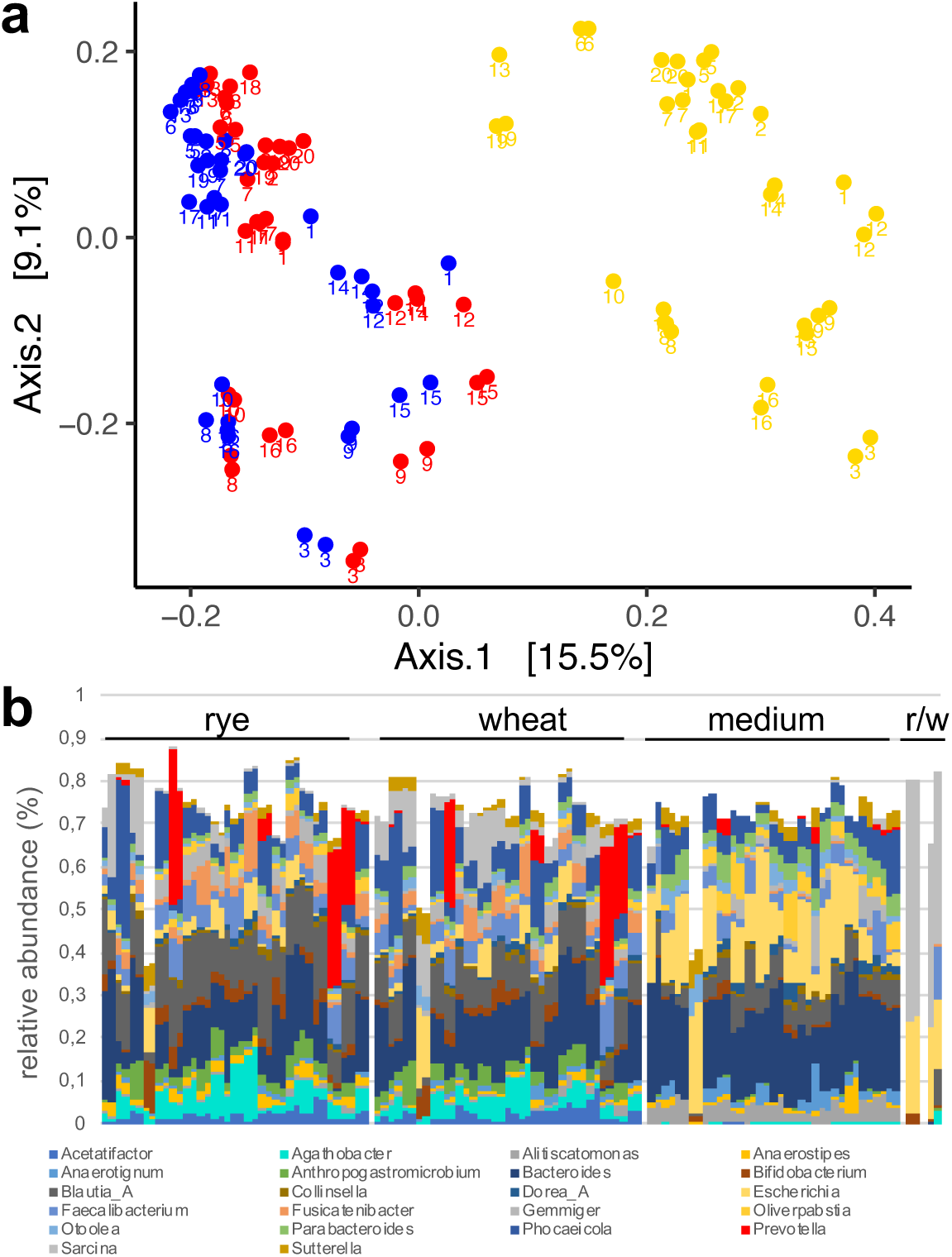
Metric dimensional scaling analysis of cultures grown on rye (blue), wheat (red) and with baseline medium only (gold) derived from 20 individuals (a). In panel b taxonomic composition on the genus level for individual substrate groups are shown. On the very right composition of cultures only containing substrate (r; rye; w: wheat) without fecal inoculum are given.

## Notes

### Competing Interest Statement

The authors have declared no competing interest.

